# Do EEG models fit users or actions? Zero-calibration BCI and limits of cross-user transfer

**DOI:** 10.64898/2026.11.27.765080

**Authors:** Tejas Bharadwaj, Unnat Parsana, Guhan Venkat

## Abstract

Brain–computer interfaces (BCIs) that decode imagined movement work with a fixed set of commands, and adding a command usually requires new labelled data from every user. A stronger goal is zero-calibration onboarding: a new action, learned once from existing users, should work for a new user who has given no labelled data. We define this problem as an onboarding curve, *A*(*n*): the area under the ROC curve (AUROC) for the new action in a held-out user, as a function of the number *n* of new-action trials pooled from other users. We measure it under a pre-specified protocol on four public motor-imagery datasets, from coarse vocabularies (hands, feet, tongue) to fine ones (individual fingers; arm and hand movements). Frozen EEG foundation models encode who the user is far better than what the user imagines. A linear probe identifies the user from single trials with 88–99% accuracy in LaBraM, CBraMod and REVE embeddings, and this identity code survives BitFit, LoRA and full fine-tuning. On five-finger imagery, no frozen foundation model recognises a newly added finger in an unseen user above 0.60 AUROC. Their cross-user signal is 16–51% of the user’s own cross-session signal, against 60–84% on coarse vocabularies. We trace this failure to four causes and build a decoder with one component per cause: a task-trained encoder with an 8–32 Hz and a 0.3–3 Hz pathway, a prototype readout, and label-free per-user re-centering. On five-finger imagery the decoder reaches *A*(512) = 0.705 in unseen users, against 0.613 for its pre-specified comparator (+0.092, 95% CI [0.073, 0.106]; 8/8 users; *p* = 0.0078). It beats every frozen foundation model in every user. Removing the 0.3–3 Hz pathway costs 0.080 AUROC, which points to slow activity as a carrier of finger information that transfers between people. Large pretrained EEG representations are dominated by individual signatures. Calibration-free fine control becomes reachable when decoders are built against that structure.

## 1 Introduction

A motor brain–computer interface (BCI) maps brain activity during imagined or attempted movement to commands [1]. Most non-invasive motor BCIs use EEG and a small command set, such as left hand, right hand, feet and tongue [2]. Two costs limit them in practice. First, EEG varies strongly between people and between sessions of the same person [3], so each user usually records labelled calibration data before use. Second, the command set is fixed when the decoder is trained. To add a command, every user must record labelled trials of it.

Many methods reduce the first cost. Subject-independent classifiers [4], zero-training approaches that reuse earlier sessions [5], Riemannian re-centering [6] and Euclidean alignment [7] all move a decoder between users with little or no labelled data from the new user [8–10]. These methods assume that the new user performs the actions the decoder was trained on. The second cost, growing the vocabulary, has received less attention. It matters most for fine control. Individual fingers or distinct arm and hand movements engage nearby cortical territory and produce weak, overlapping EEG signatures [11, 12], so collecting enough labelled trials per user for each new fine command is expensive.

We study the strongest version of the problem. A decoder is trained on a group of users for *K* actions. A new action, *K*+1, is then added from *n* trials pooled from those same users. The decoder must recognise the new action in a held-out user who has given no labelled data at all. We call this *zero-calibration onboarding*. It requires two things at once: transfer across users and extension to a class the encoder never saw. It is related to few-shot and class-incremental learning [13, 14], but the new examples come from other people.

EEG foundation models are the natural candidates for this task. Models such as LaBraM [15], CBraMod [16], REVE [17] and LUNA [18] are pretrained by masked reconstruction on thousands of hours of EEG from thousands of people. The aim is a general-purpose representation that transfers across tasks and subjects. Recent benchmarks report that frozen foundation-model features often transfer poorly under linear probing and that small specialist models trained from scratch remain competitive [19, 20]. These benchmarks measure accuracy. They do not ask what the frozen representation encodes instead. EEG carries stable individual signatures strong enough for biometric identification [21]. A representation learned without labels may therefore spend much of its capacity on who is recording rather than on what they imagine.

This paper makes four contributions:

1. We define zero-calibration onboarding as an onboarding curve *A*(*n*) and evaluate it under a pre-specified, leave-one-user-out protocol on four motor-imagery vocabularies, from coarse (BCI Competition IV-2a, Kaya HaLT) to fine (Kaya five fingers, Ofner arm and hand).
2. We show that frozen EEG foundation models fit users more than actions. They identify the user from single trials with up to 99% accuracy, and the identity code survives every adaptation level from frozen to fully fine-tuned. Their cross-user signal collapses on fine vocabularies.
3. We identify four causes of the failure on fingers and build a decoder with one component per cause.
4. On five-finger imagery the decoder recognises a newly added finger in unseen users at 0.705 AUROC, ahead of its pre-specified comparator and of every frozen foundation model in every user. Ablation attributes most of the gain to a 0.3–3 Hz pathway, which points to slow activity as a carrier of finger information that transfers between people.

## 2 Materials and methods

### 2.1 Problem definition: the onboarding curve

Let a decoder *f* = ℎ ◦ *g* consist of an encoder *g*, which maps an EEG trial to an embedding, and a head ℎ, which maps an embedding to class scores. The decoder is trained on a set of users *U* for *K* base actions. Action *K*+1 is then added by refitting the head on all base-action trials of *U* plus a random set *S_n_* of *n* trials of action *K*+1, also drawn from *U*. We call the refitted head ℎ*_n_*. The encoder *g* never sees action *K*+1.

For a held-out user *u* ∉ *U*, let *A_u_*_,*c*_ (*n*) be the AUROC with which ℎ*_n_* ◦ *g* separates trials of the new action *c* from all of *u*’s other trials, using the predicted probability of class *c* as the score. It is averaged over five random draws of *S_n_*. The onboarding curve is

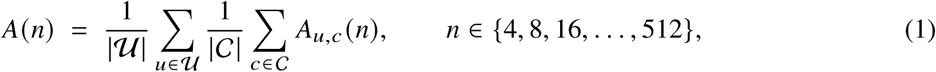

where *U* is the set of users and every class *c* ∈ C takes the role of the new action in turn. Users are held out one at a time (leave-one-user-out), so each user is in *U* once as the test user and in *U* for every other fold.

AUROC is 0.5 at chance and 1 for perfect separation. We report *A*(512) as the primary endpoint. The full curve shows the cost at smaller *n*.

*Zero calibration* means that no labelled trial of the held-out user enters training. The held-out user’s *unlabelled* trials may be used. In this study they are used for Euclidean alignment (Section 2.3) and, in our decoder, for embedding re-centering (Section 2.9). No label of *u* is used at any step. Figure 1a summarises the protocol.

**Figure 1:**
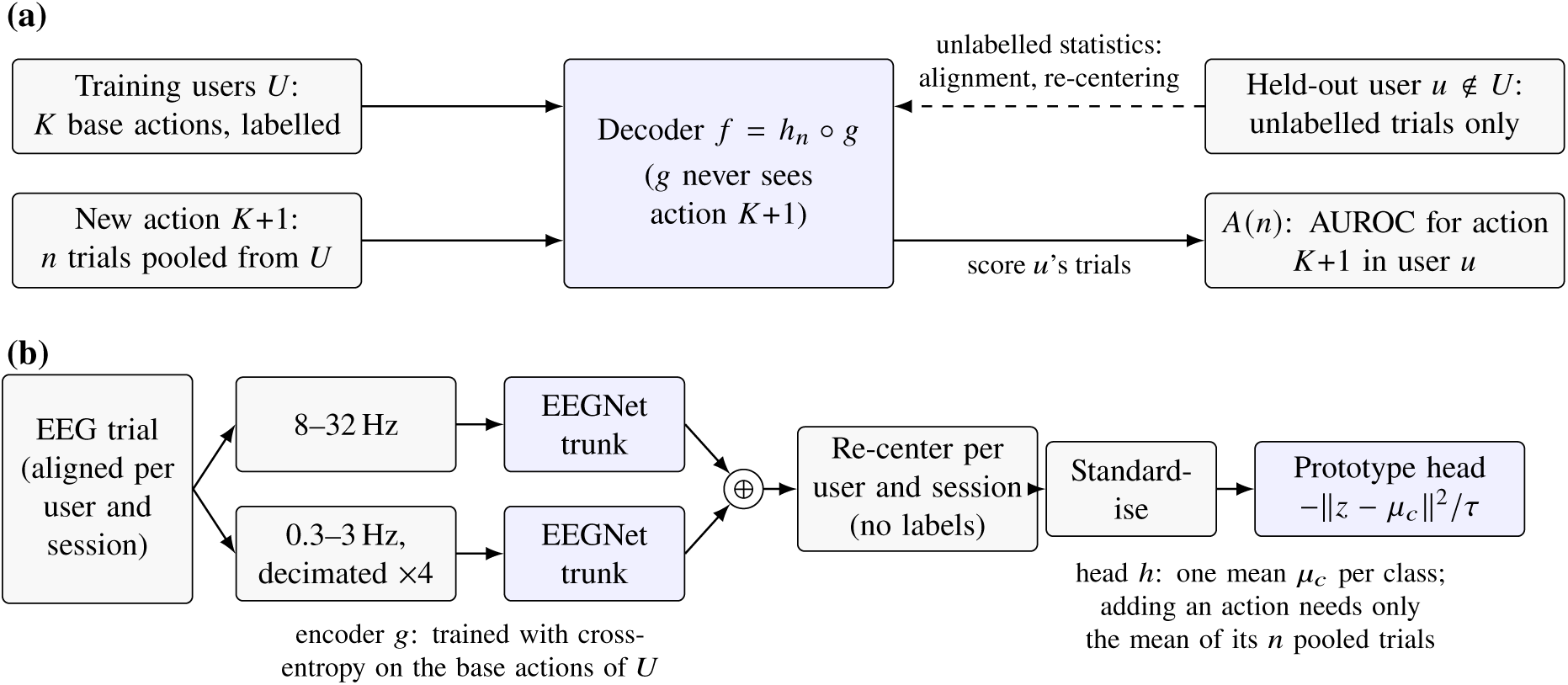
Zero-calibration onboarding and the decoder. **(a)** The decoder is trained on users *U* for *K* base actions. Action *K*+1 is added from *n* trials pooled from *U*. The decoder is then scored on a held-out user *u* who contributes only unlabelled trials. Every class takes the role of the new action in turn, and every user takes the role of *u* in turn. **(b)** The decoder. Each trial is filtered into an 8–32 Hz and a 0.3–3 Hz copy. Each copy passes through its own EEGNet trunk. The concatenated embedding is re-centered per user and session from unlabelled trials, standardised, and scored by distance to class prototypes. Blue boxes mark learned components.

### 2.2 Datasets

We used only public, de-identified EEG recorded under the original studies’ ethics approvals. No new data were collected. Four datasets served as test vocabularies (Table 1). Two further datasets, PhysioNet EEG Motor Movement/Imagery [22, 23] and OpenBMI [24], served only for development. All design decisions for the decoder were made on them. The only result we report from them is the adaptation analysis of Section 2.7, which uses PhysioNet-MI.

**Table 1:** Test vocabularies. Users and trials are those that entered the analysis. Window: epoch used by the decoder and EEGNet-emb, relative to the cue or marker.

| Dataset | Classes | Users | Trials | Channels | Window |
|---|---|---|---|---|---|
| BCI-IV-2a [25] | left hand, right hand, feet, tongue (4, coarse) | 9 | 5,184 | 22 | 4 s |
| Kaya HaLT [26] | left hand, right hand, left leg, right leg, tongue (5, coarse) | 12 | 23,081 | 19 | 1 s |
| Kaya five fingers (5F) [26] | thumb, index, middle, ring, little (5, fine) | 8 | 18,071 | 19 | 1 s |
| Ofner upper limb [12] | elbow flexion, elbow extension, supination, pronation, hand close, hand open (6, fine) | 15 | 5,400 | 61 | 3 s |

*BCI-IV-2a* contains two sessions per user (training and evaluation). We used the 2–6 s interval after the cue. *Kaya HaLT and 5F* were recorded with 22 channels [26]. We removed the two ear references and the data-acquisition channel (A1, A2, X5), leaving 19 scalp channels, and renamed T3/T4/T5/T6 to T7/T8/P7/P8. Trials were cut from 0 to 1 s after the marker onset. In HaLT we excluded the passive (no-imagery) state, leaving five imagery classes. Each recording file was treated as one session (19 files for 5F, 29 for HaLT). We analysed eight 5F users and twelve HaLT users. *Ofner* contains imagined movements of the right arm recorded in ten runs per user. We used the imagined-movement condition and the 0–3 s interval after the cue. Raw files were checked against published MD5 checksums (figshare manifest for Kaya; Zenodo record for Ofner) or, for BCI-IV-2a, against an independent download.

### 2.3 Preprocessing and alignment

Data were band-pass filtered with MNE-Python’s default FIR filter [27], resampled, and epoched, with MOABB [28] for BCI-IV-2a, PhysioNet and Ofner and a dedicated loader for Kaya. Band and sampling rate depend on the representation (Section 2.4). After epoching, each user’s trials were whitened by Euclidean alignment [7], separately for each recording session. The reference covariance of a session is the mean trial covariance, shrunk with the Ledoit–Wolf estimator [29] and, if needed, further shrunk to a condition number of at most 10^6^. Each trial is multiplied by the inverse square root of this reference. Alignment uses no labels. For the held-out user it uses only that user’s unlabelled trials.

### 2.4 Representations

We compared six representations, each followed by the same onboarding head (Section 2.5).

#### Frozen foundation models

LaBraM-Base [15], CBraMod [16], REVE-Base [17] and LUNA-Base [18] were used at their released weights without any update. Inputs were band-passed at 0.3–40 Hz and resampled to 200 Hz, as the models expect. The trial embedding is the flattened set of final-layer tokens. For Ofner, LaBraM received only the electrodes named in its position vocabulary. LUNA was evaluated on five fingers only.

#### Classical features

Trial covariance matrices were estimated with the oracle approximating shrinkage estimator [30] and projected to the Riemannian tangent space [31] (pyRiemann). The band was 8–32 Hz for BCI-IV-2a, HaLT and 5F and 0.3–3 Hz for Ofner, following the low-frequency analysis of the original Ofner study [12].

#### EEGNet-emb

An EEGNet [32] (braindecode implementation [33]; *F*_1_=8, *D*=2) was trained from scratch in every fold on the training users’ base-class trials only, with cross-entropy, Adam (learning rate 10^−3^), batch size 64 and 100 epochs. The embedding is the input to EEGNet’s final layer. Inputs were 8–32 Hz at 128 Hz (0.3–40 Hz for Ofner). Because one network is trained per (held-out user, new class) pair, EEGNet-emb never sees the held-out user or the new class. It is the task-supervised comparator for our decoder and was fixed as the primary comparator before testing.

### 2.5 Onboarding protocol and endpoints

For every representation, test vocabulary, held-out user *u* and new class *c*, the onboarding head is a logistic regression on standardised features (scikit-learn [34]; L2 penalty, *C*=1, up to 2,000 iterations). It is fitted on all base-class trials of the training users plus *n* trials of class *c* drawn without replacement from the training users, for *n* ∈ {4, 8, 16, 32, 64, 128, 256, 512} and for all available trials. Each *n* is drawn five times (seeds 0–4). The head is unweighted in the primary analysis. A class-balanced variant was run as a secondary analysis. *A_u_*_,*c*_ (*n*) is the one-vs-rest AUROC of the predicted probability of class *c* over all of *u*’s trials (Eq. 1).

As a secondary endpoint we computed an *onboarding cost*: the smallest *n* at which the mean recall of the new class reaches 80% of its recall when all available new-class trials are used. A fold enters this analysis only if its full-data recall exceeds chance (one-sided binomial test against 1/(*K*+1), *p* < 0.05). A crossing counts only if the false-alarm rate for the new class at that *n* stays at or below 1/(*K*+1). Folds that never cross are censored.

### 2.6 Identity probes

To measure how much of a representation encodes the user, we trained a linear probe (standardised features, logistic regression) to predict user identity from single trials. *Within-session identity* is the 5-fold stratified cross-validated accuracy over the training users’ trials in each leave-one-user-out fold, averaged over folds. Trials from the same session can fall into both training and test folds. Chance is 1/|*U*| (0.143 for five fingers, with seven training users). *Cross-session identity* trains the probe on each user’s first session and tests it on later sessions, so a session-specific artefact cannot explain it.

Our decoder removes each session’s mean embedding. After this re-centering every user’s mean embedding is zero, so a linear probe has no mean difference to exploit and cannot measure identity. For the decoder we therefore used a nonlinear probe: a multilayer perceptron (one hidden layer of 256 units, AdamW, learning rate 10^−3^, 100 epochs) trained on a stratified half of each training user’s trials and tested on the other half, with re-centering statistics estimated on the first half only. Before use, this probe was validated on synthetic embeddings with planted session offsets and a planted per-user code. It had to detect the planted code at more than 1.5 times chance in every fold and stay below that level when no code was planted.

### 2.7 Adaptation analysis

To test whether fine-tuning removes the identity code, we used PhysioNet-MI (users 1–30; four imagined classes: left fist, right fist, both fists, both feet). In each of 30 leave-one-user-out folds, a backbone plus a linear classification head was trained on the 29 training users for 50 epochs (batch size 64, weight decay 0.01). We compared, for CBraMod, five adaptation levels: frozen backbone (head only), BitFit [35] (bias terms), LoRA [36] on the attention query and value projections at two learning rates (10^−3^ and 10^−4^; rank 4, *α* equal to rank), and full fine-tuning. For REVE we compared frozen, BitFit and LoRA (rank 12, learning rate 10^−3^). BitFit and LoRA each trained less than 1% of the backbone’s parameters. EEGNet trained from scratch on the same folds served as reference. We report balanced accuracy on the held-out user and the identity probe (Section 2.6) on the adapted model’s embeddings of the training users’ trials. Here chance is 25% for accuracy and 1/29 = 3.4% for identity.

### 2.8 Cross-user versus within-user signal

To ask how much of a user’s own decodable signal transfers from other users, we compared two AUROCs for every user *u* and class *c*. The cross-user AUROC 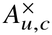 is *A_u_*_,*c*_ (512) from the onboarding protocol. The within-user AUROC 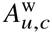 comes from a (*K*+1)-way logistic decoder trained on *u*’s own first session and tested on *u*’s later sessions. For BCI-IV-2a the split is training versus evaluation session. For Ofner, which has one session, it is runs 1–5 versus runs 6–10. We kept only cells whose within-user AUROC passed a one-sided permutation test (1,000 label permutations, *p* < 0.05), so that the ratio has a meaningful denominator, and defined

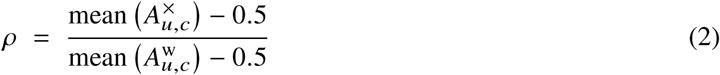

over the retained cells. *ρ* = 1 means that other users’ data are as informative about *u* as *u*’s own earlier session. Confidence intervals come from 2,000 bootstrap resamples of users.

### 2.9 The decoder

The decoder has three components, one for each cause identified in Section 3.4 (Figure 1b).

#### Task-trained two-band encoder

Each trial is band-passed twice, at 8–32 Hz (the sensorimotor *μ* and *β* bands [37]) and at 0.3–3 Hz, resampled to 128 Hz and aligned separately per band. The slow copy is decimated by 4 to 32 Hz. Each copy feeds its own EEGNet trunk (*F*_1_=8, *D*=2, temporal kernel length 64, dropout 0.25) with the classification layer removed. The two trunk outputs are concatenated into one embedding. Each band is scaled by the inverse of its standard deviation over the training trials. The encoder is trained with cross-entropy through a linear layer on the base-class trials of the training users only (Adam, learning rate 10^−3^, no weight decay, batch size 64, 100 epochs). As for EEGNet-emb, one encoder is trained per (held-out user, new class) pair, and the code asserts that neither the held-out user nor the new class enters its training set.

#### Label-free re-centering

Each embedding is centered by subtracting the mean embedding of its user and session, computed from all of that session’s trials without labels. For the held-out user this uses only unlabelled trials.

#### Prototype readout

After standardisation, each class is represented by the mean of its training embeddings (its prototype) [13, 14]. The score for class *c* is − ‖*z* − *μ_c_* ‖^2^/*τ*, where *τ* is the mean squared distance of training embeddings to their own prototype, and class probabilities are the softmax of these scores. Adding a class requires only the mean of its *n* pooled trials, so the new class does not compete with the base classes for decision boundary, as it does in a softmax classifier trained on imbalanced data [38].

Table 2 lists the full configuration. It was fixed and committed, with a hash of the encoder and head configuration, before any test vocabulary was run. The ablations remove one component at a time: the 0.3–3 Hz trunk (8–32 Hz trunk only), re-centering, or the prototype head (replaced by the logistic head of Section 2.5).

**Table 2:** Decoder configuration. , fixed before testing.

| Component | Setting |
|---|---|
| Input bands | 8–32 Hz and 0.3–3 Hz; 128 Hz; slow band decimated ×4 |
| Alignment | Euclidean alignment per user and session, Ledoit–Wolf shrinkage, per band |
| Trunks | 2 × EEGNet (*F*<sub>1</sub>=8, *D*=2, kernel 64, dropout 0.25), classification layer removed |
| Encoder training | cross-entropy on base classes of training users; Adam, lr 10<sup>−3</sup>, batch 64, 100 epochs; seed = fold index |
| Re-centering | subtract per-(user, session) mean embedding; unlabelled |
| Head | standardisation; nearest-prototype, squared Euclidean distance;<br>temperature = mean within-class squared distance |
| Onboarding | prototype of new class = mean of *n* pooled trials; *n* ∈ {4, . . . , 512}, 5 draws per *n* |

### 2.10 Pre-specification and statistics

The protocol, endpoints, comparators, test and correction were recorded in a version-controlled pre-registration before the decoder was run on any test vocabulary. The primary hypothesis was that the decoder’s *A*(512) exceeds that of EEGNet-emb. For each vocabulary we computed the per-user difference Δ*_u_* = 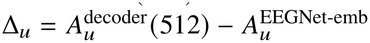, each term averaged over new classes, and tested mean(Δ*_u_*) with an exact two-sided sign-flip test over all 2^|*u*|^ sign assignments [39]. With eight users the smallest attainable *p* is 2/2^8^ = 0.0078. We corrected across the four vocabularies with the Holm procedure [40] at *α* = 0.05 (step-down thresholds 0.0125, 0.0167, 0.025, 0.05). Confidence intervals for mean(Δ*_u_*) are 95% percentile bootstrap intervals over users (2,000 resamples). All other comparisons, including those with the frozen foundation models, the ablations and the analyses of Sections 2.6–2.8, are descriptive and carry no hypothesis test.

### 2.11 Software and reproducibility

Analyses used Python 3.12 with PyTorch [41], braindecode [33], MNE-Python [27], MOABB [28], pyRie-mann and scikit-learn [34], with every dependency pinned in a lockfile. Training used deterministic algorithms where available and a fixed seed per fold. Every result row records the code commit that produced it, and the analysis scripts refuse to write results from an uncommitted state. Backbone encoding, adapters and the decoder ran on a CUDA GPU. All other analyses ran on CPU.

## 3 Results

### 3.1 Frozen foundation models encode users more than actions

On five-finger imagery, the representations that encode the user most strongly onboard a new finger worst (Table 3). In LaBraM, CBraMod and REVE embeddings, a linear probe identifies the user from a single trial with 88–99% accuracy (chance 14%). Yet none of these models recognises a newly added finger in an unseen user above 0.60 AUROC. The two task-trained representations show the opposite pattern. EEGNet-emb and our decoder carry the weakest identity code (0.46 and 0.49) and reach the highest AUROC (0.613 and 0.705). Identity does not order the frozen models among themselves: REVE has both the strongest identity code and the best frozen AUROC.

**Table 3:** Identity versus onboarding on five-finger imagery. (8 users). Identity: within-session linear identity probe (chance 0.143). *A*(512): new-finger AUROC in the held-out user with the linear onboarding head. With an MLP head, REVE reaches 0.632 and CBraMod 0.627.

| Representation | Type | Identity probe | *A*(512) |
|---|---|---|---|
| REVE | frozen foundation model | 0.994 | 0.594 |
| CBraMod | frozen foundation model | 0.976 | 0.582 |
| LaBraM | frozen foundation model | 0.884 | 0.511 |
| LUNA | frozen foundation model | 0.716 | 0.545 |
| Classical (tangent space) | hand-designed | 0.536 | 0.558 |
| EEGNet-emb | task-trained | 0.461 | 0.613 |
| **Our decoder** | task-trained | 0.49<sup>†</sup> | **0.705** |

<sup>†</sup>Nonlinear split-half probe after re-centering (Section 2.6); a linear probe is uninformative after re-centering.

The identity code is not a session artefact. A probe trained on each user’s first session and tested on later sessions still identifies the user with 0.748 accuracy from REVE embeddings, against 0.202 from classical features (chance 0.143).

A nonlinear readout does not close the gap. With a multilayer-perceptron head instead of the linear head, *A*(512) rises to 0.632 for REVE and 0.627 for CBraMod. This is above EEGNet-emb’s linear value (0.613) but 0.07–0.08 below the decoder.

### 3.2 Adaptation does not remove the identity code

Fine-tuning improves accuracy but leaves identity intact (Figure 2a, Table 4). On PhysioNet-MI, CBraMod’s balanced accuracy rises from 40.9% frozen to 53.2% after full fine-tuning, and its identity probe stays between 93% and 100% at every level. REVE stays at 47–51% accuracy and 95–99% identity at all three levels tested. No adaptation level beats EEGNet trained from scratch (53.5%), which identifies the user with only 16% accuracy. Fine-tuning a pretrained backbone thus buys no accuracy over a small supervised network, while it keeps an identity code that the supervised network does not acquire.

**Figure 2:**
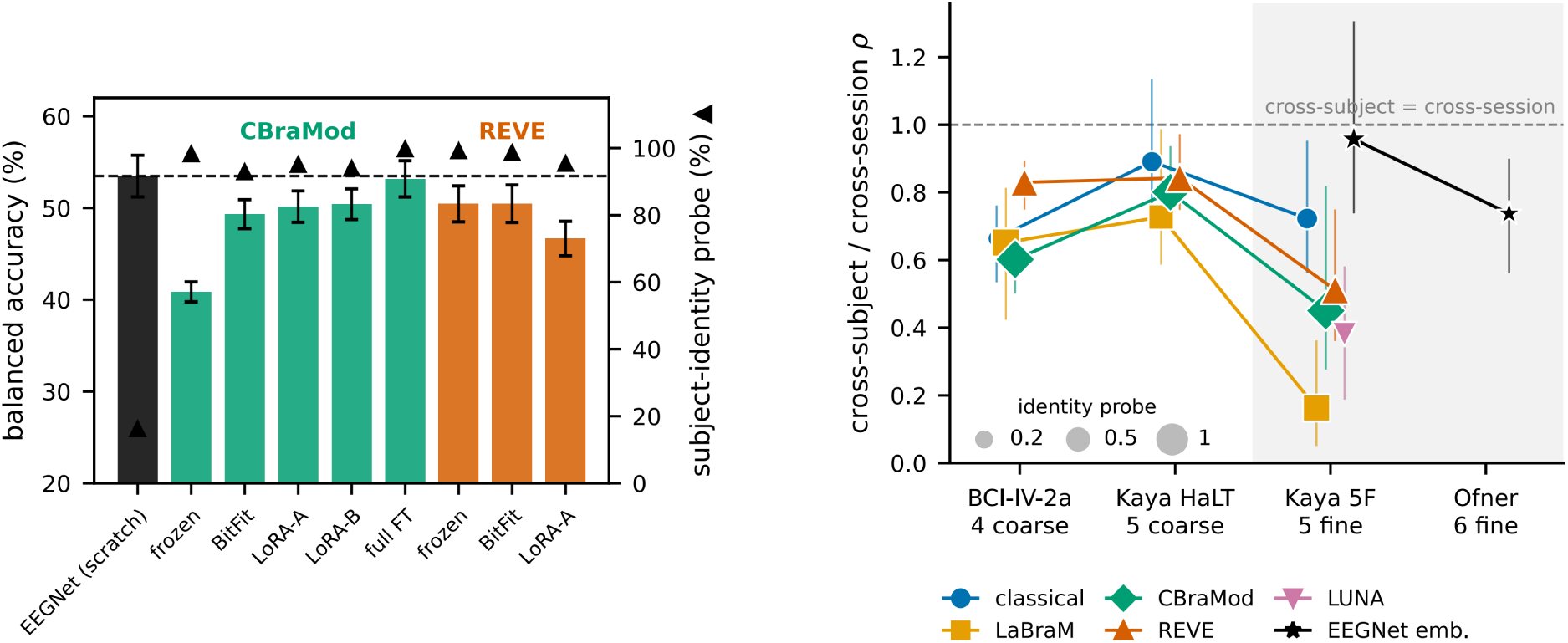
Foundation models fit users, not actions. **(a)** PhysioNet-MI, 4 classes, 30 users: balanced accuracy on the held-out user (bars) and identity-probe accuracy (triangles, right axis) from frozen to fully fine-tuned. LoRA-A and LoRA-B: learning rates 10^−3^ and 10^−4^. Dashed line: EEGNet from scratch. No adaptation level beats EEGNet from scratch (53.5% vs. 53.2% for fully fine-tuned CBraMod), which identifies the user at only 16%. **(b)** Cross-user signal as a fraction of the user’s own cross-session signal (*ρ*, Section 2.8), from coarse to fine vocabularies (fine shaded). Marker area is proportional to the identity probe. Error bars: 95% bootstrap intervals over users. Descriptive.

**Table 4:** Adaptation ladder on PhysioNet-MI. (30 users, 4 classes). Balanced accuracy on the held-out user (chance 25%) and identity probe on the adapted embeddings (chance 3.4%). Values as plotted in Figure 2a.

| Model | Adaptation | Balanced accuracy (%) | Identity (%) |
|---|---|---|---|
| EEGNet | trained from scratch | 53.5 | 16.1 |
| CBraMod | frozen | 40.9 | 98.2 |
|  | BitFit | 49.3 | 92.9 |
|  | LoRA, lr 10<sup>−3</sup> | 50.1 | 95.0 |
|  | LoRA, lr 10<sup>−4</sup> | 50.4 | 93.9 |
|  | full fine-tuning | 53.2 | 99.7 |
| REVE | frozen | 50.5 | 99.1 |
|  | BitFit | 50.5 | 98.6 |
|  | LoRA, lr 10<sup>−3</sup> | 46.7 | 95.4 |

### 3.3 Cross-user transfer collapses on fine vocabularies

For frozen foundation models, the share of a user’s own signal that transfers from other users falls sharply from coarse to fine vocabularies (Figure 2b, Table 5). On BCI-IV-2a and HaLT, their cross-user AUROC is 60–84% of the within-user cross-session AUROC. On five fingers it is 16–51%, with LaBraM lowest (0.16). Classical features hold up better on fingers (*ρ* = 0.72). The task-trained EEGNet-emb keeps nearly all of its within-user signal across users on fingers (*ρ* = 0.96) and 0.74 on Ofner. The loss on fine vocabularies is therefore not inherent to fine imagery. It is largest for frozen pretrained representations.

**Table 5:**
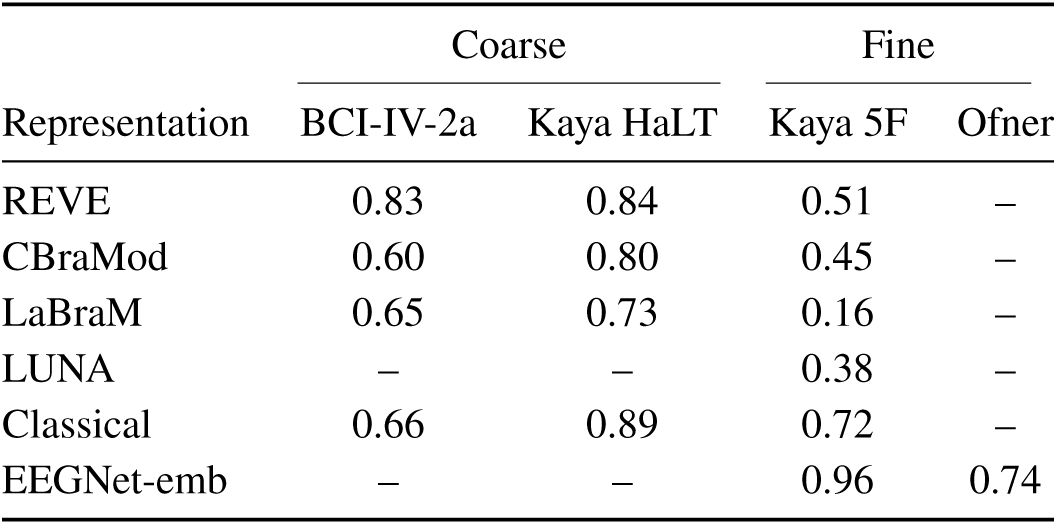
Cross-user over within-user signal, *ρ*, by representation and vocabulary. *ρ* = 1: other users’ data are as informative as the user’s own earlier session. Values as plotted in Figure 2b; –: not estimated.

| Representation | Coarse |  | Fine |  |
|---|---|---|---|---|
|  | BCI-IV-2a | Kaya HaLT | Kaya 5F | Ofner |
| REVE | 0.83 | 0.84 | 0.51 | – |
| CBraMod | 0.60 | 0.80 | 0.45 | – |
| LaBraM | 0.65 | 0.73 | 0.16 | – |
| LUNA | – | – | 0.38 | – |
| Classical | 0.66 | 0.89 | 0.72 | – |
| EEGNet-emb | – | – | 0.96 | 0.74 |

These ratios should be read against a low denominator. Fine imagery is weak even within one person: a user’s own decoder trained on one session and tested on later ones reaches 22–32% balanced accuracy on five fingers (chance 20%), against 39–66% on coarse BCI-IV-2a (chance 25%).

### 3.4 Each failure has a cause, and each cause sets a design choice

The analyses above point to four causes of the failure on fingers (Table 6). We built the decoder with one component per cause.

**Table 6:** From findings to design. Each finding maps to one component of the decoder.

| Finding | Evidence | Component of the decoder |
|---|---|---|
| Task training beats frozen pretraining on the finger vocabulary | EEGNet-emb 0.613 > every frozen model (Table 3); *ρ* = 0.96 (Table 5) | Encoder trained on the task (EEGNet trunks [32]), not frozen |
| Slow EEG activity carries part of the finger signal | Within-user audit; low-frequency upper-limb decoding [12] | Second trunk on 0.3–3 Hz beside 8–32 Hz |
| Softmax starves a newly added class | Base:new imbalance 6.75:1 to 35:1 at *n* = 512 | Prototype readout [13] in place of softmax |
| User and session shift the embedding | Identity recovery at 0.88–0.99 (Table 3) | Label-free re-centering per user and session |

#### Frozen pretraining loses to task training on fine vocabularies

EEGNet-emb, trained on only seven users’ base fingers, beats every frozen foundation model under the pre-specified linear head (0.613 vs. at most 0.594) and keeps the most within-user signal across users (*ρ* = 0.96). The decoder’s encoder is therefore trained on the task rather than frozen.

#### Part of the finger signal is slow

A within-user audit showed decodable finger information below the 8– 32 Hz band that motor-imagery decoders usually use, consistent with low-frequency time-domain decoding of upper-limb movements [12, 42]. The decoder adds a second trunk on 0.3–3 Hz.

#### A softmax head starves a newly added class

At *n* = 512 the base classes outnumber the new class by 6.75:1 (BCI-IV-2a) to 35:1 (HaLT) in the head’s training set. A softmax classifier fitted on such data biases its decisions against the minority class [38]. The decoder replaces it with a prototype readout, in which the new class is one mean and needs no decision boundary fitted against the others.

#### Users and sessions shift the embedding

Identity is recoverable at 0.88–0.99 from frozen embeddings, which means each user and session occupies its own region of embedding space. The decoder removes each user-session mean without labels.

### 3.5 The decoder onboards a new finger without calibration

Tested once per vocabulary against EEGNet-emb, the decoder raises *A*(512) on five fingers from 0.613 to 0.705 (Δ = +0.092, 95% CI [0.073, 0.106]; positive in 8/8 users; exact sign-flip *p* = 0.0078, below the Holm threshold of 0.0125). This *p* is the smallest attainable with eight users. Per-user gains over EEGNet-emb range from +0.03 to +0.11 (Figure 3b).

**Figure 3:**
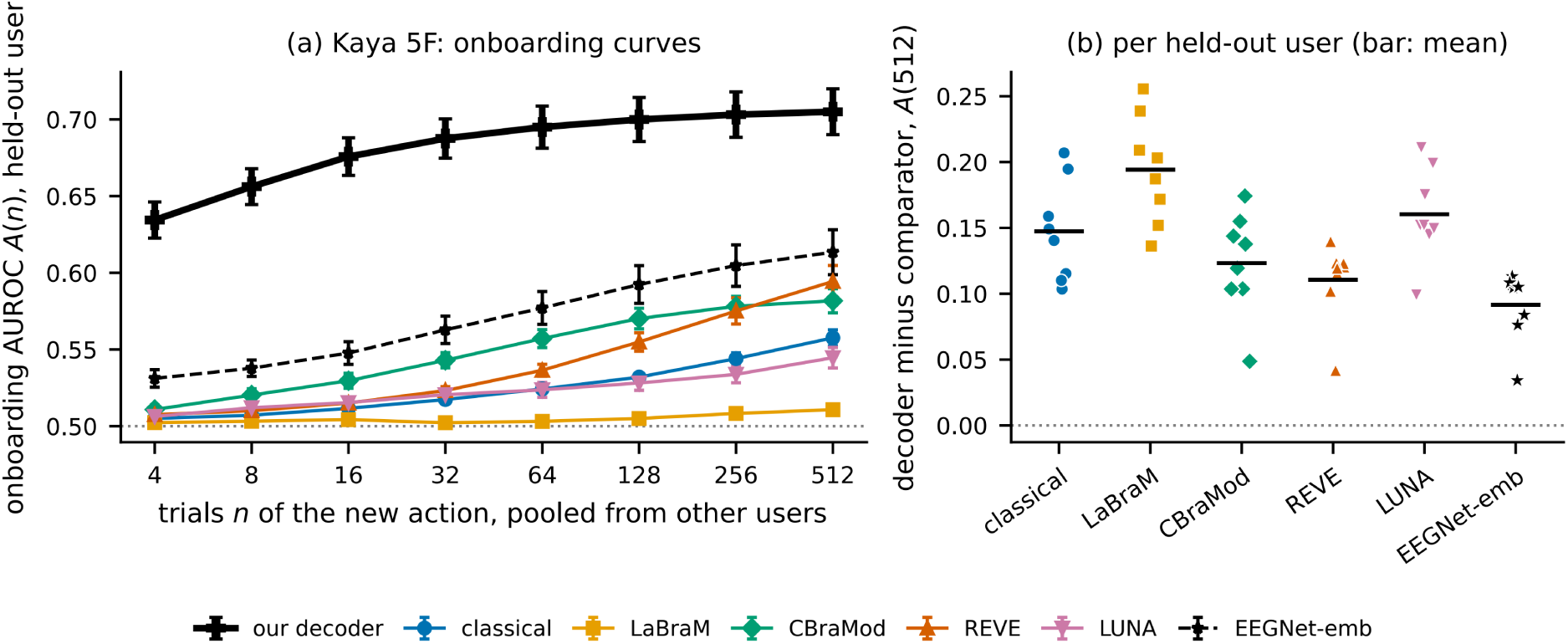
Zero-calibration onboarding of a new finger. (Kaya five fingers, 8 users). **(a)** Onboarding curve *A*(*n*): AUROC for the new finger in a held-out user against the number *n* of new-finger trials pooled from other users (mean ± s.e.m. over users). Dotted line: chance. **(b)** Per-user gain of the decoder over each comparator at *n* = 512; bar: mean. The decoder is ahead in every user against every comparator.

The decoder also leads every frozen foundation model in every user (descriptive): by +0.111 over REVE, +0.123 over CBraMod, +0.147 over classical features, +0.160 over LUNA and +0.194 over LaBraM, each in 8/8 users (Figure 3b).

The advantage holds along the whole onboarding curve (Figure 3a, Table 7). With only *n* = 4 pooled trials of the new finger, the decoder reaches 0.634. This exceeds every frozen model at *n* = 512 and EEGNet-emb at *n* = 512. The decoder is within 0.01 of its final value by *n* = 64 (0.695), whereas REVE and EEGNet-emb still climb at *n* = 512. LaBraM stays near chance at every *n* (0.502–0.511).

**Table 7:** Onboarding curves on five fingers. : *A*(*n*) for each representation, mean over 8 held-out users and 5 new-finger roles. Values as plotted in Figure 3a.

|  | Pooled new-finger trials *n* |  |  |  |  |  |  |  |
|---|---|---|---|---|---|---|---|---|
| Representation | 4 | 8 | 16 | 32 | 64 | 128 | 256 | 512 |
| **Our decoder** | **0.634** | **0.656** | **0.676** | **0.688** | **0.695** | **0.700** | **0.703** | **0.705** |
| EEGNet-emb | 0.531 | 0.538 | 0.548 | 0.563 | 0.577 | 0.592 | 0.605 | 0.613 |
| REVE | 0.508 | 0.510 | 0.515 | 0.523 | 0.537 | 0.555 | 0.575 | 0.594 |
| CBraMod | 0.511 | 0.520 | 0.530 | 0.543 | 0.557 | 0.570 | 0.578 | 0.582 |
| Classical | 0.505 | 0.507 | 0.512 | 0.517 | 0.524 | 0.532 | 0.544 | 0.558 |
| LUNA | 0.506 | 0.512 | 0.515 | 0.521 | 0.524 | 0.528 | 0.534 | 0.545 |
| LaBraM | 0.502 | 0.503 | 0.504 | 0.502 | 0.503 | 0.505 | 0.508 | 0.511 |

For the secondary onboarding-cost endpoint, re-centering raised the number of measurable five-finger folds with a finite onboarding cost (80% criterion) from 27 to 34 of 38.

### 3.6 The slow pathway carries most of the gain

Removing one component at a time shows that the 0.3–3 Hz trunk carries most of the decoder’s advantage (Table 8). Without it, *A*(512) on five fingers falls by 0.080, from 0.705 to about 0.625, which is close to EEGNet-emb (0.613). The same removal costs 0.074 on HaLT but only 0.014 on BCI-IV-2a. Recentering leaves *A*(512) unchanged (−0.001 to −0.003), although it increases the number of folds with a finite onboarding cost (above). Replacing the prototype head with the logistic head changes *A*(512) by −0.003 on fingers and *improves* it on the coarse vocabularies (+0.055 on HaLT, +0.030 on BCI-IV-2a). In AUROC terms, the gain on fingers therefore comes from the task-trained two-band encoder, chiefly its slow trunk.

**Table 8:** Ablations. : change in *A*(512) when one component is removed. Prototype head: replaced by the logistic head. Descriptive.

| Component removed | Kaya 5F (fine) | Kaya HaLT (coarse) | BCI-IV-2a (coarse) |
|---|---|---|---|
| 0.3–3 Hz trunk | −0.080 | −0.074 | −0.014 |
| Re-centering | −0.001 | −0.003 | −0.001 |
| Prototype head | −0.003 | +0.055 | +0.030 |

### 3.7 Coarse and arm–hand vocabularies

The decoder’s advantage is specific to fingers. The pre-specified test was significant after Holm correction on one of four vocabularies, five fingers. On the coarse vocabularies the decoder reached *A*(512) = 0.772 on HaLT and 0.690 on BCI-IV-2a, which ties or trails the best frozen foundation model. On Ofner’s arm and hand movements it did not differ significantly from EEGNet-emb.

## 4 Discussion

We asked whether EEG representations fit users or actions, and what that means for adding a command to a BCI without calibrating it on the new user. Frozen foundation models fit users. Their embeddings identify the person from single trials almost perfectly, fine-tuning does not remove this code, and on fine motor vocabularies little of the action signal transfers from other users. A decoder designed against this structure, trained on the task, re-centered per user and reading out by prototypes, recognises a newly added finger in an unseen user at 0.705 AUROC, ahead of every comparator in every user.

### Why pretrained EEG representations encode identity

One explanation is that masked reconstruction rewards whatever explains the most variance in the signal. In EEG, stable individual factors such as head geometry, skull conductivity, electrode contact and resting rhythms contribute large, consistent variance [3, 21]. A representation that predicts missing EEG well must therefore model these factors, while the variance that separates one imagined finger from another is small. Our results agree with this account. The identity code is strong in every foundation model, survives a session change, and persists through every adaptation level, whereas a network trained on the motor task from scratch identifies users far less well (16% on PhysioNet-MI, chance 3.4%; 0.46 on five fingers, chance 0.14). This also explains why benchmarks find frozen features weak under linear probing [19, 20]: the linear head must find the action signal inside a space organised largely by identity.

### Slow activity as a transferable finger code

The 0.3–3 Hz trunk accounts for most of the decoder’s gain on fingers, and its removal costs more on the fine vocabulary than on the coarse BCI-IV-2a. Low-frequency, time-domain EEG carries information about upper-limb movements [12, 42], and slow movement-related cortical potentials precede and accompany voluntary movement [43]. Our finding adds that such slow activity may carry finger information that is shared across people, whereas the 8–32 Hz band that motor-imagery decoders usually use [2, 37] transfers less. Two other sources could also produce class-specific slow activity and are not excluded here. The first is cue-evoked potentials: in the Kaya paradigm the target action is shown by a visual cue, and our 1-s window starts at the marker. The second is ocular activity, such as eye movements toward cue positions, which concentrates at low frequencies [44]. Testing the slow pathway on windows that exclude the cue response, with ocular regression, is the most important next step.

### What the prototype head and re-centering do

In AUROC terms neither component helped on fingers, and the prototype head was worse than the logistic head on coarse vocabularies. This is consistent with what AUROC measures. AUROC is threshold-free, so it ignores the decision bias that class imbalance induces in a softmax head [38]. That bias shows up in recall and false alarms, which our onboarding-cost endpoint captures, and there re-centering raised the number of folds with a finite onboarding cost from 27 to 34 of 38. The two components should therefore be judged on decision-level endpoints, and our evidence for them is weaker than for the encoder.

### Implications for foundation models and for privacy

Two practical implications follow. First, an evaluation of an EEG representation for BCI should report an identity probe and a cross-user onboarding curve, not only within-dataset accuracy. A high score on a coarse benchmark says little about fine vocabularies, where cross-user transfer of frozen features collapsed. Second, pretrained EEG embeddings identify their users. Embeddings shared or stored in place of raw EEG should be treated as identifying personal data [45, 46].

### Limitations

The five-finger dataset has eight users, so the primary test reached the smallest *p*-value attainable with eight users and the estimates have wide uncertainty [47]. The decoder was significant on one of four vocabularies. On coarse vocabularies it ties or trails the best frozen foundation model, so it is not a general replacement. The slow trunk doubles the encoder’s size, so its effect is not separated from added capacity. Cue-evoked and ocular contributions to the slow signal are untested (see above). Alignment and re-centering use a batch of the new user’s unlabelled trials from the same session, including the trials being scored. An online system would need running estimates from a short unlabelled period, which we did not test. All data are offline recordings from healthy participants. The adaptation analysis covers two foundation models on one development dataset. Some values in Tables 4, 5 and 7 are reported as plotted in the corresponding figures.

## Conclusion

Large pretrained EEG representations are dominated by individual signatures rather than by the shared motor code. For BCI, fine-grained, calibration-free control becomes reachable once decoders are built against that structure: trained on the task, aware of the user and session, and open to slow activity.

## Data availability

All datasets are public. BCI Competition IV 2a: https://www.bbci.de/competition/iv/. Kaya et al. HaLT and 5F: https://doi.org/10.6084/m9.figshare.c.3917698.v1. Ofner et al. upper limb: https://doi.org/10.5281/zenodo.834976. PhysioNet EEG Motor Movement/Imagery (development): https://doi.org/10.13026/C28G6P. OpenBMI (development): https://doi.org/10.5524/100542. Pretrained foundation-model weights are from their public releases.

## Code availability

Code is available from the authors on reasonable request.

## Ethics statement

This study used only publicly available, de-identified EEG data collected under the ethics approvals of the original studies. No new data were collected from human participants.

